# Forest canopy cover composition and landscape influence over bryophytes communities in *Nothofagus* forests of southern Patagonia

**DOI:** 10.1101/2020.04.27.063826

**Authors:** Mónica DR. Toro Manríquez, Víctor Ardiles, Álvaro Promis, Alejandro Huertas Herrera, Rosina Soler, María V. Lencinas, Guillermo J. Martínez Pastur

**Author notes:** These authors contributed equally to this work (Mónica DR. Toro Manríquez, Víctor Ardiles). These authors also contributed equally to this work (Álvaro Promis, Alejandro Huertas Herrera, Rosina Soler, María V. Lencinas, Guillermo J. Martínez Pastur).

## Abstract

Understanding the influence environmental drivers on understory vegetation is important for conservation efforts under climate change. Bryophytes are one of the most diverse groups in temperate forests but also the least known. In addition, the environmental drivers (e.g., forest structure, microclimate, soil conditions or substrate) influencing over bryophyte community among *Nothofagus* forest types are poorly known. The aim of this study was to evaluate the influence of forest canopy-layer composition on the structure (cover) and the composition (richness and diversity) of bryophyte communities (mosses and liverworts) in two contrast landscape types (coast and mountain) in southern Patagonia. Three natural *Nothofagus* forest types (pure deciduous, pure evergreen, and mixed deciduous-evergreen) in two landscapes (coast < 100 m.a.s.l.; mountain > 400 m.a.s.l.) were selected (N = 60 plots). In each forest plot, we established one linear transect (10 m length) to measure bryophyte cover (point-intercept method). The data were evaluated using ANOVAs, Chi-square test and multivariate analyses. The mosses were mostly austral-antarctic origin, and the liverworts were all endemics. The principal substrates for the bryophytes development in the forest floor were litter and decaying woods. Moreover, many bryophytes species act as a substrate for natural tree regeneration. The forest structure was the main driver of bryophytes community in the coast landscape, while the slope was the principal driver of bryophytes in the mountain landscape. These differences were mainly explained for the microclimate into the forests (e.g., soil moisture and air temperature), and for the regional climate in the landscapes (e.g., air temperature and soil conditions). Notably, the mixed forest, mainly in the coast, presented exclusive species that were not present in the deciduous and evergreen pure forests. The conservation efforts should include management considerations both the stand and landscape levels based on the potential climate-change impact over bryophyte communities.

## Introduction

The bryophytes are worldwide-distributed plants in different terrestrial ecosystems [1]. In forests, the bryophytes play crucial ecological functions (e.g., carbon cycle, nutrients cycle, water balance) [2, 3], and an important role for seed germination, seedling growth and forest regeneration [4–6]. Such role affects the local biodiversity [7, 8], and bryophytes composition is related to the canopy cover, the microclimate and site conditions in the stands (e.g., forest structure, humidity, light intensity, substrates, soil moisture, soil pH) [9, 10]. The bryophytes have a broader distribution and a longer altitudinal gradient than vascular plants (e.g., woody plants, ferns) [6]. Many species show wide transcontinental ranges and sometimes the occurrence of bryophyte species may be a vestige of the moss flora prior to the rupture of Laurasia and Gondwana [11]. Exogenous effects such as narrow ecological requirements (e.g., climate, habitat) can explain this discrepancy [12].

In southern Patagonia, native forest occupying large area are dominated by *Nothofagus* tree species: the deciduous *Nothofagus antarctica* (G.Forst.) Oerst. and *Nothofagus pumilio* (Poepp et. Endl) Krasser; and the evergreen *Nothofagus betuloides* (Mirb.) Oerst. These tree species grow together and can form deciduous-evergreen mixed forests, being the *N. pumilio* and *N. betuloides* mixed-forest the most representative [13, 14]. Here, the understory is mainly composed of forbs, grasses and few woody species, where deciduous and mixed forests are similar in species composition, but mountain forests sustain highest herb-layer diversity and more particular species than forests in the coast (i.e., strong influence of landscape type) [15]. However, bryophytes appear to have been neglected in many ecological studies of Patagonian terrestrial ecosystems and these coupled effects (forest type and landscape type) have not been examined for bryophytes.

Currently, forest research on the diversity, richness and distribution of bryophytes had been increasing because of their role in ecosystem functioning and the interest in facing climate change [6, 16, 17]. While most species have been described in southern Patagonia highlighting the dependence on wetter forests [18, 19], knowledge about bryophytes distribution, diversity patterns in different ecosystems (e.g., forests, grasslands, peatlands), substrate requirements (e.g., litter, decaying wood) and their response to ecological drivers along environmental gradients is poor. Few studies focused on community ecology of bryophytes in *Nothofagus* forests e.g., [20, 21]. Among them, Lencinas et al. [20] described high variability in the ground-bryophyte communities of deciduous *Nothofagus* forests related to forest structure heterogeneity and different microhabitat or substrates conditions. Therefore, it is not clear how different environmental drivers (e.g., canopy cover and composition, temperature, substrate) effect on bryophyte communities distribution in native forests, including pure and mixed stands in different landscapes of southern Patagonia. Comprehensive knowledge of the diversity and its spatial patterns is required for effective planning and management at both stand and landscape scale.

The aims of this study were to evaluate the influence of forest canopy-layer composition on bryophyte richness, cover and diversity at two contrast landscape in Tierra del Fuego, and determine how bryophyte community vary according to the environmental drivers at stands and landscape levels. The following specific questions were asked: (1) What are the ecological drivers influencing bryophyte composition in the understory? (2) How these ecological drivers vary from pure to mixed forests and how this affects the bryophyte composition? (3) How does the influence of these environmental drivers vary in two contrasting environments within the distribution of mixed forests? (4) Are the substrate or the microclimate more relevant? Complementary, the global distribution patterns and substrates of bryophytes will be described in relation to the mentioned factors (forest type and landscape type); and the relationship between bryophyte plants (mosses and liverworts) with the presence of forest seedlings will be analysed. This study might contribute to identify strategies and opportunities for the conservation of bryophyte species and to generate basic knowledge about the potential effects of climate change in temperate forest landscapes.

## Methods

### Study area

This study was conducted in old-growth *Nothofagus* forest stands (>250 years old) located in the southwest of Tierra del Fuego Island, Argentina (Fig 1), without harvesting or cattle grazing during the last 50 years. Three *Nothofagus* forest types were considered, according to their canopy-layer composition: (i) Np = pure deciduous forests with >80% of *N. pumilio* canopy cover; (ii) Nb = pure evergreen forests with >80% *N. betuloides* canopy cover; and (iii) M = mixed *N. pumilio* - *N. betuloides* forests, with similar proportion of deciduous and evergreen species in the canopy. Likewise, two landscapes were selected, where these three forest types occurred: (i) marine coast close to the Beagle Channel, and (ii) mountain areas toward the inner island. The coast landscape was placed within the Tierra del Fuego National Park, where the altitude varied between 50 and 100 m.a.s.l., the mean annual air temperature was 4.3°C, and the annual precipitation recorded reached 756 mm year^−1^ with abundant snowfalls during winter. The stands in the mountain landscape were placed in Garibaldi Pass (450 m.a.s.l.), within the Andes Mountain range, where the annual temperature was 3.1°C, and the annual precipitation recorded was 788 mm year^−1^. The soil type is loam textured, with massive granular structures, low usable water capacity and moderate to slow internal and external drainage with a thick soil layer [22]. Canopy composition in these forests has an impact on soil properties, but also most soil properties are strongly influenced by landscape (i.e., mountain soils are wetter and richer in N and P than soils in the coast) [14]. In addition, litterfall (quantity and quality) which determines nutrient cycling in soils also differ among forest types and landscape types [14].

**Fig 1.**
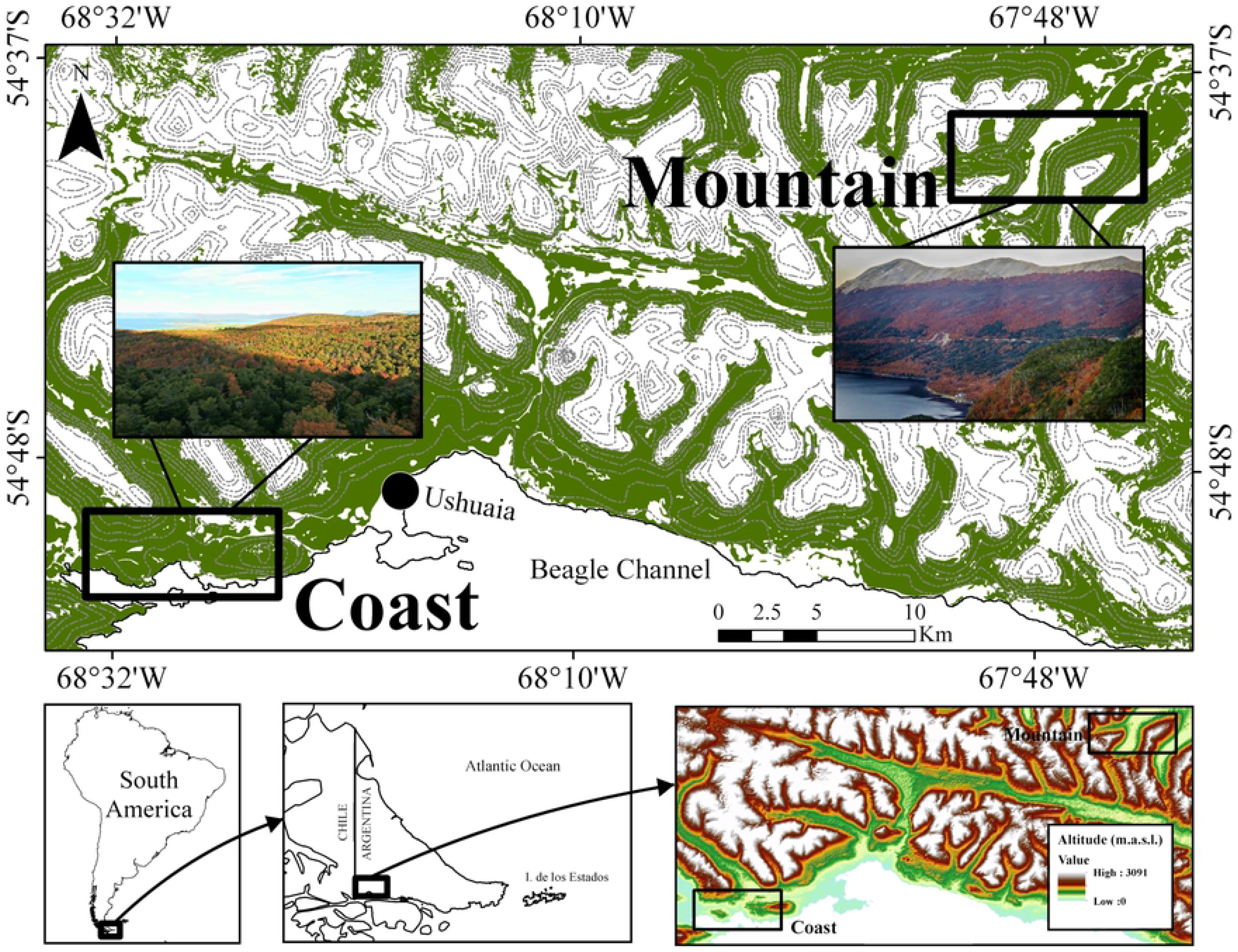
Location of the study area in southwest Argentinean part of Tierra del Fuego and altitude (m.a.s.l.). Rectangles indicate sampling landscape types: Coast (the coast of the Beagle Channel) and Mountain (the Andean Mountains region). In green the *Nothofagus* forests. Photos next to rectangles are an example of the analyzed forest landscapes (photo by A. Huertas Herrera and J.M. Cellini).

### Forest structure, microclimate and forest floor

For this study, three forest types (Np, Nb and M) and the two landscape types (coast and mountain) were selected. The forest structure was characterized by angle-count sampling plots (N = 10 × 3 forest types × 2 landscape types = 60 plots), following Bitterlich [23]. We used a Criterion RD-1000 (Laser Technology, USA) with a variable basal area factor between 6 and 7, to calculate tree basal area (BA, m^2^ ha^−1^). Additionally, dominant height (DH, m) was measured using a TruPulse 200 laser clinometer and distance rangefinder (Laser Technology, USA) by the averaging height of the three taller trees per plot. In addition, the species and diameter at breast height (DBH, cm) were recorded for each tree. At each plot, hemispherical photos of the canopy were taken. We used a 8 mm fish-eye lens (Sigma Ex-AF4, Japan) mounted on a 35 mm digital camera (Nikon D50, Japan) with a tripod and level, which was oriented to the North. The camera was set 1 m above the forest floor, which is enough to avoid registering understory or shrub cover in these austral forests and only capture tree canopy cover. The program Gap Light Analyzer v2.0 [24] was used to calculate canopy cover (CC, %), relative leaf area index (RLAI) and transmitted total radiation (TR, %) (ratio of direct plus diffuse radiation transmitted through the canopy, and the total radiation incident on a horizontal surface located above forest canopy). These data were obtained by taking one photo per plot each month along the growing season (October to March) avoiding the influence of direct sunlight.

Twelve data loggers (model H8, HOBO, USA) were used to characterize the microclimate considering forest types and landscape types, measuring soil temperature (ST, °C), air temperature (AT, °C), relative air humidity (RH, %). Finally, to measure the effective annual precipitation (PP, mm yr^−1^) through the canopy, twelve Spectrum Watermark Watchdog 425 data loggers (Spectrum, USA) were installed. All microclimatic drivers were measured between October 2014 to March 2015.

During January 2015 were measured the soil resistance to penetration and soil moisture taking five measurements per stand. The soil resistance to penetration (R, N cm^−2^) was determined by a manual penetrometer (Eijkelkamp Agrisearch Equipment, Netherlands) up to 30 cm deep, while the soil moisture (SM, %) was determined with an MP406 moisture probe (ICT, Australia) in the first 6-10 cm of soils. At the same time, five mineral soil samples from each stand (N = 60 total samples) were taken at 0-20 cm deep with a soil auger. The samples were dried in an oven at 70°C, ground in an analytical mill type Cole-Parmer (USA) and then sieved (2 mm). The soil pH was also determined at the Agroforestry Resources Laboratory (CADIC) using a pH-meter Orion (USA). The slope or terrain (S, %) was calculated for each site with a Suunto clinometer (Finland).

### Sampling design and bryophyte composition

In forest plots, we established one linear transect (10 m length) (N = 60 plots). For the measurements, we employed the point-intercept method [25] completing 50 intercept points per transect (every 20 cm) to collect and record bryophyte species considering two groups: mosses and liverworts. Specimens of hornworts group (*Anthocerotophya* division) were not found in the transects. With these data, we calculated the species richness (moss richness and liverwort richness) and cover in percentage (moss cover and liverwort cover) for each transect. We also recorded other forest floor covers, such as bare soil (BS, %) which refers to forest soil usually covered by litter but without vegetation, woody debris cover (Ds, %) that included branches and trunks >3 cm in diameter, vascular plants (VC, %) that include ferns, monocots and dicots, and the lichens cover (L, %).

The fieldwork was conducted during summer (2015) which is the most appropriated period for vegetation surveys in *Nothofagus* forests at this latitude (within the growing season) [20]. For species determination, we followed the methodology proposed by Ardiles et al. [26] and Ardiles and Peñaloza [27]. We carried out microscopic analysis of specimens for structural details, confirming and/or complementing the taxonomic information, at the Botanical Area of the Museo Nacional de Historia Natural (Santiago, Chile), being able to observe in detail specific parts of the gametophytes and sporophytes. We used the nomenclature proposed by Müller [28] for moss species and that of Hässel de Menéndez and Rubies [29] for liverworts. To establish the global distribution patterns (GDP) of each species, we followed the proposal of León et al. [30], which is essentially an adaptation of the patterns proposed by Seki [31] and Villagrán et al. [32]. According to them, we classified the species as: D = disjunt with South America, South Africa and Europe; E = endemic; PAN = pantropical-type *Podocarpus*; A = austral-antarctic. We also analysed the occurrence frequency (OF, %) and the main taxonomic groups (Ms = mosses and Li = liverworts) (S1 Table). Finally, the substrate (understood as microhabitat) where the specimens were growing was registered at each sampling point in the transect: litter (LT), decaying wood (DW), bare soil (BS), stone (St), and epiphytic on branches and/or bark in the forest floor (EP) whose development was prior to falling to forest floor. At each point, the presence of tree seedlings growing on bryophytes was also recorded. We then calculated the frequency of tree regeneration on bryophytes for each forest and landscape types.

### Data analyses

We first calculated bryophyte richness and relative cover and calculated Shannon-Wiener diversity (H’) and Pielou evenness indices (J’) [33]. Then, two-way ANOVAs considering the forest types (Np, M, Nb) and landscape types (COA = coast, MOU = mountain) as main factors, were used to analyze forest structure (BA, DH, DBH, CC, RLAI), microclimate (GR, SM, PP, ST, AT, RH), soil and forest floor conditions (S, pH, R, BS, Ds, VC, TL), mosses richness, liverworts richness, mosses cover, liverworts cover, and diversity indexes (H’, J’). Likewise, forest structure (BA, DH, DBH) differences between overstory tree species were compared in mixed forests, also using two-way ANOVAs, where tree species (*N. pumilio* and *N. betuloides*) and landscape types (coast and mountain) were considered as main factors for these analyses. For VC and L were ln(Y + 1) transformed for the analyses to accomplish statistical assumptions, but non-transformed mean values are shown in the Tables. The comparisons of mean values were conducted using Tukey honestly significant difference test (p <0.05). Statistically significant interactions were plotted for better interpretation of results. Considering the microhabitat analyses and the tree seedlings growing on bryophytes, we also compared the treatments using contingency tables and the Chi-Square test, considering comparisons at forest type and landscape type: CNp= deciduous *Nothofagus pumilio* forests in the coasts, CM= mixed *N. pumilio* and *N. betuloides* forests in the coasts, CNb = evergreen *N. betuloides* forests in the coasts, MNp = deciduous *N. pumilio* forests in the mountains, MM = mixed *N. pumilio* and *N. betuloides* forests in the mountains, MNb = pure evergreen *N. betuloides* forests in the mountains. Statistical analyses were conducted using Statgraphics (Statistical Graphics Corp., USA) and IBM SPSS software.

Multivariate methods were also conducted (using the species cover data), as complementary analyses: (i) Indicator species analysis (IndVal) for bryophytes species composition, comparing among forest types and between landscape types. The maximum value of the IndVal between the groups (forest types and landscape types, separately) was evaluated to determine its statistical significance (p <0.05) using a Monte Carlo permutation test (number of runs 4999) [34]. We follow the criteria used by [21], which considers an indicator species of non-vascular plants with an IndVal ≥25 and the values of p <0.05. (ii) Canonical correspondence analyses (CCA) based on species cover data (mosses and liverworts) were used to analyze the relationships between bryophyte community and environmental drivers. Firstly, Pearson correlation was calculated for the environmental drivers (p <0.05), assigned to them a value between −1 and 1, (above 0.5 or below −0.5 indicate the existence of correlation). Likewise, Monte-Carlo method with 499 permutations was employed to test the significance of each axis in CCA. All multivariate analyses were performed using PC-ORD [35].

## Results

### Forest structure, forest floor cover and microclimate

The forest structure showed significant differences among forest types and landscapes types, except for canopy cover (%) (Table 1). The basal area was higher in evergreen forests (84.5 m^2^ ha^−1^) than in mixed forests (77.1 m^2^ ha^−1^) and deciduous forests (66.5 m^2^ ha^−1^). Likewise, the basal area was higher in the mountain (80.7 m^2^ ha^−1^) than in the coast (71.4 m^2^ ha^−1^). Dominant height and diameter at breast height presented differences along a gradient: deciduous forests > mixed forests > evergreen forests. RLAI was higher and similar in deciduous and mixed forests than in evergreen forests. Likewise, RLAI was significantly higher on the coast (2.5) than in the mountain (2.3). Climate drivers showed significant differences among forest types (e.g., soil temperature and relative air humidity) and landscape types (e.g., soil moisture and relative air humidity). The soil temperature was higher in deciduous (6.2°C) > evergreen (5.7°C) > mixed forests (4.3°C), while relative air humidity was higher in mixed (62%) > evergreen (55%) > deciduous forests (39%). In addition, landscape type showed differences for soil moisture, which were higher in the mountain than in the coast, and relative air humidity was higher in the coast than in the mountain. The soil and forest floor conditions also showed significant differences among forest types (e.g., slope, pH, bare soil, vascular plants cover) and between landscape types (e.g., bare soil, resistance to penetration, lichen cover). The slope and bare soil presented higher values in evergreen (14% and 31%, respectively) and mixed forests (13% and 39%, respectively) compared to deciduous forests (7% and 16 %, respectively). Contrary, pH was higher for deciduous forest > mixed forests > evergreen forests. Beside this, resistance to penetration, bare soil and lichen cover were higher on the coast than in mountain. There were significant interactions for basal area, relative air humidity, lichen cover, pH, and resistance to penetration (Table 1; Fig 2). On the other hand, basal area, dominant height and diameter at breast height did not differ significantly between *N. pumilio* and *N. betuloides* species in mixed forests (S2 Table). Neither of these differences was found between landscapes type, only in diameter at breast height (coast > mountain). Interactions were not significant either (S2 Table). The interactions in basal area occurred for all forest types had similar values, which of deciduous forests had significantly lower basal area than mixed and evergreen forests in the mountain. The basal area in the coast and mountain landscapes differed only in mixed forests (Fig 2). The relative air humidity was similar in the deciduous forests in landscape types. However, the mixed and evergreen forests, presented higher relative air humidity in the coast (Fig 2). Interaction in lichen cover occurred by significantly higher cover in coast than in mountain in mixed and evergreen forests (Fig 2), but not in the deciduous. Statistical differences were not detected among forest types for each landscape type. In pH, a significant interaction was detected due to all forest types had significantly values on the coast, but mixed and evergreen forests had similar pH values in the mountains (Fig 2). The resistance to penetration was similar in all forest types in the mountain, but on the coast it was significantly higher in deciduous than in mixed and evergreen forests. However, the resistance to penetration was significantly higher in the coast than in the mountain (Fig 2).

**Table 1.** Two-way ANOVAs evaluating the effect of forest types (Np = deciduous forests with >80% *Nothofagus pumilio* canopy cover, M = mixed forests of *N. pumilio* and *N. betuloides*, Nb = evergreen forests with >80% *N. betuloides* canopy cover) and landscape type (COA = coast, MOU = mountain) over: (i) Forest structure: BA = basal area (m^2^ ha^−1^), DH = dominant height (m), DBH = diameter at breast height (cm), CC = canopy cover (%); RLAI = relative leaf area index. (ii) Climate: TR = transmitted total radiation (%), SM = soil moisture (%), PP = annual precipitation (mm yr^−1^), ST = soil temperature (°C), AT = air temperature (°C), RH= relative air humidity (%).(iii) Soil and forest floor conditions: S = slope (%), pH = pH of the upper 10 cm of the soil, R = resistance to penetration (N m^−2^), BS = bare soil cover (%), Ds = woody debris cover up to 5 cm diameter (%), VC= vascular plant cover including ferns, monocots and dicots (%), L= lichen cover (%).

| Factor |  | Forest structure |  |  |  |  | Climate |  |  |  |  |  | Soil and forest floor conditions |  |  |  |  |  |  |
| --- | --- | --- | --- | --- | --- | --- | --- | --- | --- | --- | --- | --- | --- | --- | --- | --- | --- | --- | --- |
|  |  | BA | DH | DBH | CC | RLAI | TR | SM | PP | ST | AT | RH | S | pH | R | BS | Ds | VC | L |
| Forest types | Np | 66.5 a | 21.9 b | 63.2 c | 86.1 | 2.5 b | 18.7 | 31.4 | 55.8 | 6.2 b | 6.7 | 38.7 a | 7.1 a | 4.9 c | 392.8 | 16.5 a | 14.7 | 15.2 b | 0.6 |
|  | M | 77.1 ab | 17.6 a | 47.9 b | 86.2 | 2.5 b | 18.5 | 34.9 | 51.5 | 4.3 a | 6.9 | 61.5 b | 12.7 b | 4.3 b | 288.3 | 38.8 b | 17.7 | 6.5 a | 0.9 |
|  | Nb | 84.5 b | 15.9 a | 37.5 a | 86.8 | 2.1 a | 18.1 | 39.3 | 47.8 | 5.7 b | 7.2 | 54.8 b | 12.9 b | 3.7 a | 250.5 | 31.4 b | 17.3 | 6.9 a | 0.7 |
|  | *F* | 7.12 | 12.31 | 33.35 | 0.44 | 11.12 | 0.19 | 1.47 | 0.48 | 5.59 | 0.25 | 8.78 | 4.87 | 24.52 | 2.40 | 11.51 | 1.14 | 32.9 | 0.41 |
|  | *p* | 0.002 | <0.001 | <0.001 | 0.645 | <0.001 | 0.826 | 0.240 | 0.624 | 0.010 | 0.779 | 0.001 | 0.012 | <0.001 | 0.099 | <0.001 | 0.327 | <0.001 | 0.663 |
| Landscape types | COA | 71.4 a | 18.7 | 51.7 | 86.2 | 2.5 b | 18.8 | 13.4 a | 55.2 | 5.7 | 7.3 | 58.6 b | 10.1 | 4.1 | 474.2 b | 36.6 b | 17.3 | 8.8 | 1.0 b |
|  | MOU | 80.7 b | 18.3 | 47.4 | 86.5 | 2.3 a | 18.1 | 57.0 b | 48.1 | 5.1 | 6.6 | 44.7 a | 11.7 | 4.5 | 146.8 a | 21.2 a | 15.8 | 10.3 | 0.4 a |
|  | *F* | 5.76 | 0.20 | 2.81 | 0.20 | 5.50 | 0.53 | 135.07 | 1.16 | 1.43 | 2.12 | 9.36 | 0.84 | 3.95 | 78.69 | 15.70 | 0.68 | 2.56 | 5.56 |
|  | *p* | 0.020 | 0.683 | 0.100 | 0.655 | 0.023 | 0.471 | <0.001 | 0.292 | 0.243 | 0.158 | 0.005 | 0.364 | 0.052 | <0.001 | <0.001 | 0.414 | 0.116 | 0.006 |
| Interaction | *F* | 5.71 | 1.80 | 0.75 | 2.03 | 1.98 | 0.90 | 2.84 | 3.81 | 0.30 | 0.08 | 4.02 | 1.20 | 3.87 | 3.20 | 0.32 | 0.78 | 3.50 | 5.46 |
|  | *p* | 0.006 | 0.175 | 0.476 | 0.142 | 0.148 | 0.414 | 0.067 | 0.050 | 0.746 | 0.927 | 0.031 | 0.311 | 0.027 | 0.040 | 0.730 | 0.462 | 0.050 | 0.007 |

*F*, *p* = Fisher test and probability. Different letters in each column show significant differences by Tukey test at *p* < 0.05. For VC and L were ln(Y + 1) transformed for the analyses, but non-transformed data are shown.

**Fig 2.**
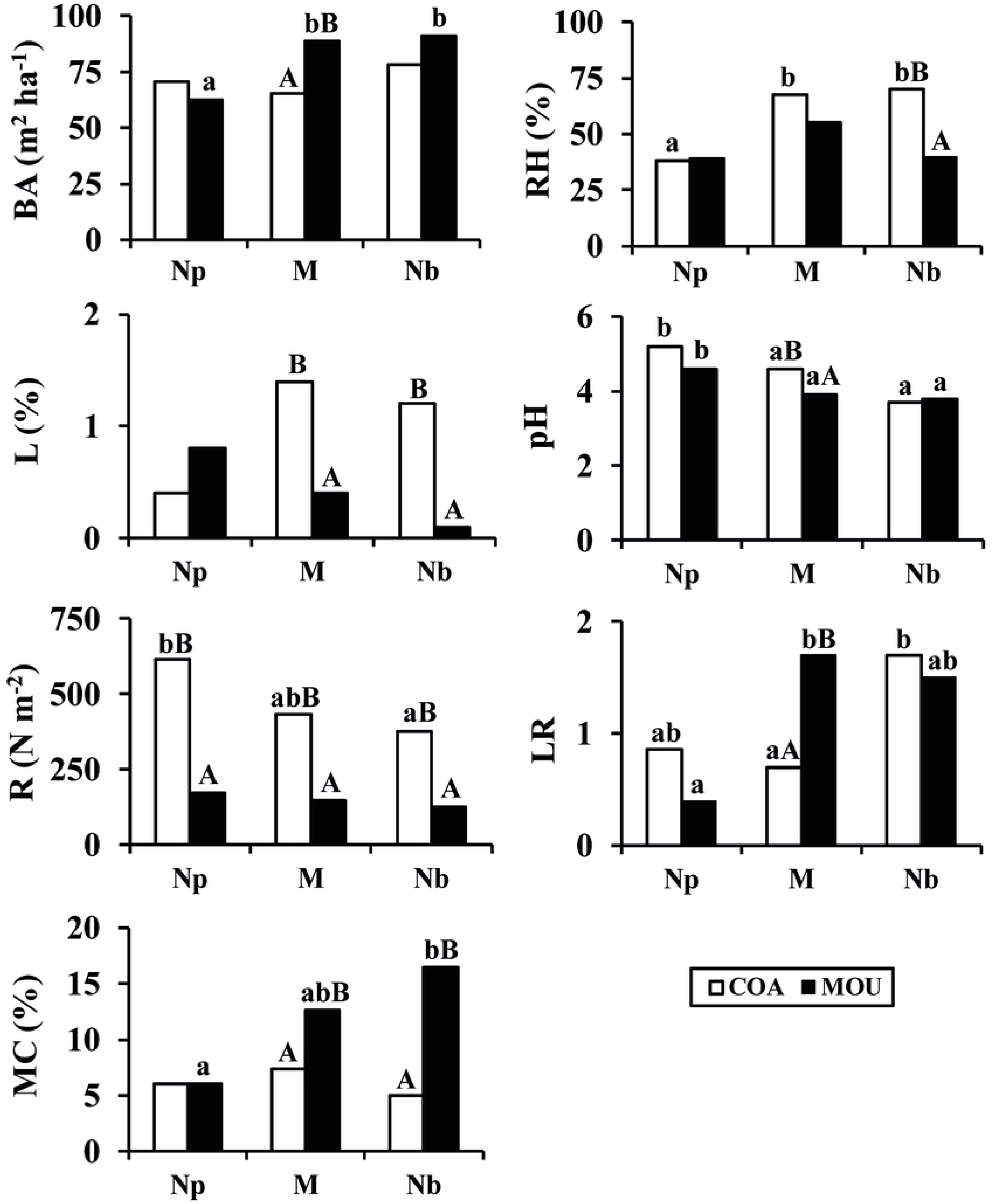
Interactions corresponding to Table 1 and Table 2. BA = basal area (m^2^ ha^−1^), RH = relative air humidity (%), L = lichen cover (%), pH = pH of the upper 10 cm of the soil, R = resistance to penetration (N m^2^), MC = mosses cover (%), LR = liverworts richness. Different letters showed significant differences by Tukey test (*p* <*0.05*). Lower cases were used for comparisons among forest types, and capital letters were used for comparisons between landscape types (COA = coast, MOU = mountain).

**Table 2.** Two-way ANOVAs evaluating the effect of forest types (Np = deciduous forests with >80% *Nothofagus pumilio* canopy cover, M = mixed forests of *N. pumilio* and *N. betuloides*, Nb = evergreen forests with >80% *N. betuloides* canopy cover) and landscape types (COA = coast, MOU = mountain). Richness and cover for mosses and liverworts, and Diversity indices: H’ = Shannon-Wiener diversity index, J’ = Pielou evenness index.

|  |  | Richness |  | Cover (%) |  | Diversity indices |  |
| --- | --- | --- | --- | --- | --- | --- | --- |
| Factor | | Mosses | Liverworts | Mosses | Liverworts | $H'$ | $J'$ |
| Forest types | Np | 1.6 a | 0.6 a | 6.0 | 1.9 | 0.64 a | 0.95 b |
|  | M | 2.6 b | 1.2 ab | 10.0 | 3.7 | 1.06 b | 0.92 ab |
|  | Nb | 2.0 ab | 1.6 b | 10.7 | 5.0 | 1.05 b | 0.88 a |
|  | *F* | 3.29 | 5.18 | 2.90 | 2.73 | 3.90 | 3.22 |
|  | *p* | 0.045 | 0.009 | 0.064 | 0.075 | 0.027 | 0.048 |
| Landscape types | COA | 1.8 | 1.1 | 6.1 a | 3.1 | 0.90 | 0.94 |
|  | MOU | 2.3 | 1.2 | 11.7 b | 3.9 | 0.94 | 0.90 |
|  | *F* | 2.26 | 0.220 | 11.01 | 0.57 | 0.08 | 3.64 |
|  | *p* | 0.139 | 0.639 | 0.002 | 0.454 | 0.777 | 0.062 |
| Interaction | *F* | 0.31 | 3.48 | 3.82 | 2.61 | 1.13 | 1.73 |
|  | *p* | 0.736 | 0.038 | 0.028 | 0.083 | 0.329 | 0.188 |

*F*, *p* = Fisher test and probability. Different letters in each column show differences by Tukey test at $p < 0.05$ .

### Bryophyte characterization, richness, cover and diversity index

A total of 27 bryophytes species were recorded: 56% were mosses (15 species; *Bryophyta* division) and 44% were liverworts (12 species of liverworts; *Marchantiophyta* division, *Jurgemanniopsida* Sub-class) (S1 Table). The global distribution patterns for the recorded species showed that mosses corresponded to austral-antarctic (53%), pantropical-type *Podocarpus* (20%), and cosmopolitan, bipolar or circum- antarctic (7% for each one). The liverworts corresponded to endemic (58%), Disjunt with South America, South Africa and Europe (33%), and small proportions were undetermined (8%). The most common bryophytes were the moss *Acrocladium auriculatum*, registered principally in deciduous forests and mountain landscape (48% mean frequency among forests types, and 32% between landscape types), and the liverwort *Adelanthus lindbergianus*, mainly in the evergreen forests of the mountains (32% mean frequency among forests types, and 21% between landscape types). The lowest occurrence frequencies (1.7% in average among forest types, and 1.1% in average between landscape types) were found in the coast landscape, three of which were mosses (*Bartamia mossmaniana* and *Hymenodontopsis mnioides* in mixed forests, and *Hennediella densifolia* in deciduous forests) and two were liverworts (*Heteroscyphus intergrifolius* in deciduous forests and *Lophozia sp.* in mixed forests).

The liverworts were found in all microhabitats, and they not presented significant differences in the association among microhabitats or forest-landscape types (Chi-square = 22.9; df = 20; p = 0.289). In contrast, mosses presented significant differences for its association (Chi-square = 77.08; df = 25; p <0.001). *Acrocladium auriculatum* was the moss species with the major variety of microhabitats, it was not found only on stones. *Adelanthus lindbergianus* and *Lepidozia chordulifera* were the liverworts with the major variety of microhabitat, they were not found only on stones (S3 Table).

The most common moss species, *Acrocladium auriculatum*, was significantly associated (Chi-square = 28.9; df = 12; p = 0.004) with some microhabitats (litter, decaying wood, bare soil, and epiphytes) considering the forest type-landscape (except for CNb, where this species was not found). Another moss, *Dicranoloma robustum*, was significantly associated (Chi-square = 18.2; df = 5; p = 0.003) with two microhabitats (litter and DW) and all forest types of different landscapes. Within the most common liverworts, *Leptocyphus hiudobroanus*, was found only in the forests in the coast, and was statistically associated (Chi-square = 16.1; df = 6; p = 0.013) with litter, decaying wood, bare soil and epiphytes.

The mosses and liverworts richness showed significant differences among forest types (Table 2). Only MC presented significant differences between landscape types (higher in the mountain than in the coast). Shannon-Wiener diversity (H’) and Pielou evenness (J’) indices showed significant differences among forest types (Table 2). H’ was higher for mixed (1.06) and evergreen forests (1.05) than in deciduous forests (0.64), while J’ was higher for deciduous (0.95) and mixed forests (0.92) than in evergreen forests (0.88).

There were significant interactions for liverwort richness, mosses cover (Table 2; Fig 2). Interactions in liverwort richness were due to in coast and mountain landscapes all forest types have significant differences, but presenting low values in deciduous forests. At the same time, liverwort richness of the coast and the mountain differed only in mixed forests (Fig 2). The mosses cover was similar in deciduous forests at both landscape types. However, the mosses cover in mixed and evergreen forests was significantly higher in the mountain compared with the coast (Fig 2).

Several shared species were observed among the different forest types (33% of richness = 9 species), which were found both in deciduous and evergreen forests (Fig 3). This shared richness was higher when was compared among the different forest types: deciduous vs. mixed forests (37%, 10 shared species) and evergreen vs. mixed forests (67%, 18 shared species). The highest exclusive species was observed in evergreen forests (18%, 5 species), compared with those in deciduous (7%, 2 species) and mixed forests (4%, 1 species). Regarding the landscape types, the shared species reached 52% (14 shared species), 30% (8 exclusive species) for the coasts and 18% (5 exclusive species) for the mountains. The IndVal showed that the mosses *Dicranoloma robustum* and *Ditrichum cylindricarpum* were more frequent in evergreen forests, *Acrocladium auriculatum* in deciduous forests, and *Campylopus clavatus* in mixed forests (Table 3). Regarding landscapes, the liverwort *Leptoscyphus huidobroanus* and the moss *Dicranoloma robustum* were indicator species in coast, while the mosses *Campylopus clavatus* and *Ditrichum cylindricarpum* were indicator species in the mountain (Table 3).

**Fig 3.**
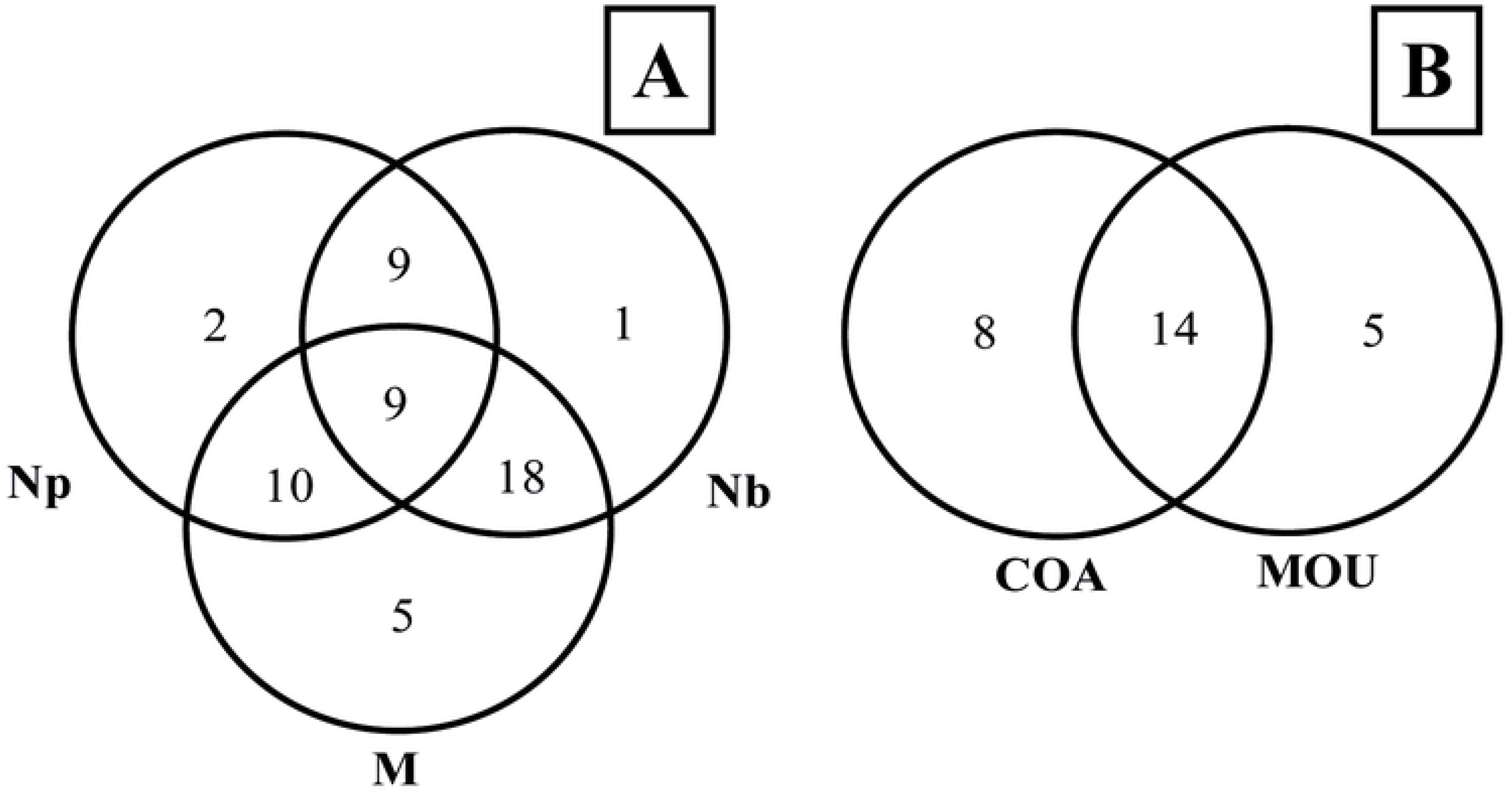
Overlapping diagrams for bryophytes richness (common and exclusive species). (A) forest types (Np = deciduous forests with >80% *Nothofagus pumilio* canopy cover, M = mixed forests of *N. pumilio* and *N. betuloides*, Nb = pure evergreen forests with >80% *N. betuloides* canopy cover), and (B) landscape types (COA = coast, MOU = mountain).

**Table 3.** Indicator species analyses for bryophyte composition at each forest type (Np = deciduous forests with >80% *Nothofagus pumilio* canopy cover, M = mixed forests of *N. pumilio* and *N. betuloides*, Nb = pure evergreen forests with >80% *N. betuloides* canopy cover) and landscape types (COA = coast, MOU = mountain).

| Forest types |  |  | Landscape types |  |
| --- | --- | --- | --- | --- |
| Np | M | Nb | COA | MOU |
| *Acrocladium auriculatum* | *Campylopus clavatus* | *Dicranoloma robustum* | *Leptoscyphus huidobroanus* | *Campylopus clavatus* |
| (38.6)* | (25.0)* | (30.0)* | (34.6)** | (25.8)** |
|  |  | *Ditrichum cylindricarpum* | *Dicranoloma robustum* | *Ditrichum cylindricarpum* |
|  |  | (28.9)* | (30.4)* | (30.4)* |

The indicator values are reported between brackets. \*= significance at $p < 0.05$ , \*\*=indicated significance at $p <$ 0.01.

The frequency of tree regeneration growing over bryophytes showed a significant association between the different liverwort (Chi square = 33.5; df = 21; p = 0.041) and mosses species (Chi square = 118.4; df = 28; p <0.001) considering the different forest types and landscapes (S4 Table). However, one exception exists for CM, where no regeneration plants were recorded over bryophytes. This association was more frequent in mountain forests, mainly in MM and MNb, where liverworts species as *Adelanthus lindbergianus* and *Lepidozia chordulifera* were highly associated with *Nothofagus* tree seedlings. On the other hand, moss species as *Acrocladium auriculatum* and *Dicranoloma robustum* were highly associated with tree seedlings in all forests types at each landscape type.

### Influence of forest structure, microclimate and floor cover on bryophyte distribution

The CCA explained 67% of variance for species-environment, with a total inertia of 7.3 and eigenvalues of 0.485 in Axis 1 and 0.366 in Axis 2 (Fig 4). The environmental drivers employed for the CCA were previously selected for its statistically significance according to Pearson correlation coefficient (Table 4; S5 Table). Axis 1 was influenced by annual precipitation and RLAI and Axis 2 was influenced by relative air humidity, air temperature and slope. When species were analyzed alone (Fig 4A), both axes showed a close correlation and association between mosses and liverworts according to microclimatic drivers. Moreover, the CCA separated the sampling plots in two main groups, defined by its landscape type (coast and mountain), with few differences between the different forests types (Fig 4B). The environmental drivers were the most related to the coasts forests (e.g., RLAI, annual precipitation, relative air humidity, air temperature), while slope influenced in the mountain forests.

**Fig 4.**
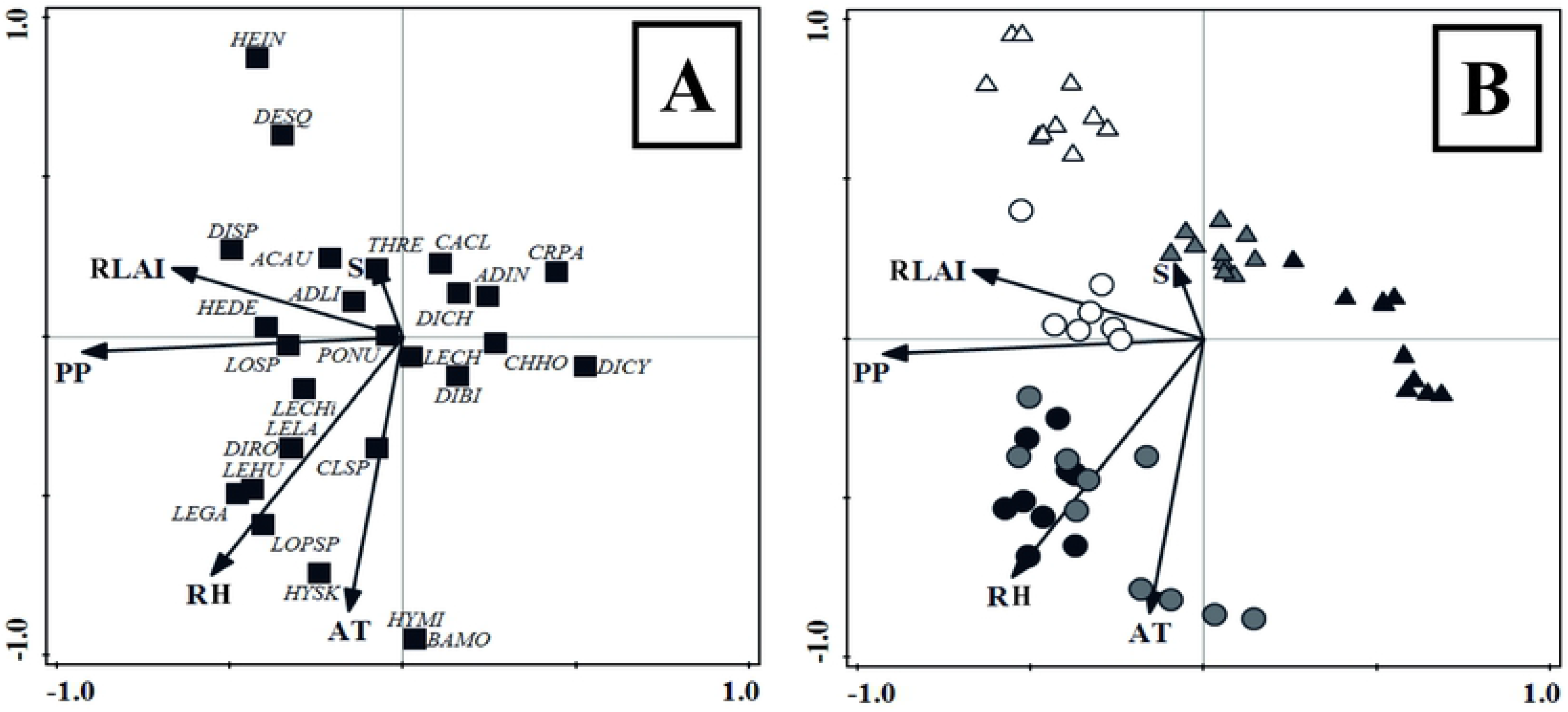
Canonical correspondence analysis based on species abundance to asses the influence of environmental drivers (RLAI = relative leaf area index, S = Slope (%), PP = annual precipitation (mm yr^−1^), RH = relative air humidity (%), AT = air temperature (°C) on (A) bryophyte species distribution (see S1 Table for species code), and (B) studied stands (deciduous *N. pumilio* forests in the coasts (CNp) and mountains (MNp), evergreen *N. betuloides* forests in the coasts (CNb) and mountains (MNb), mixed forests in the coasts (CM) and mountains (CM)).

**Table 4.** Significant drivers obtained in the Canonical Correspondence Analysis model, including explained variation, contribution, pseudo-F test, and associated probability *p* < 0.05. PP = annual precipitation (mm yr^−1^), RH = relative air humidity (%), RLAI = relative leaf area index, AT = air temperature (°C), S = slope (%).

| Drivers | Variation explained % | Contribution % | pseudo-F | *p* |
| --- | --- | --- | --- | --- |
| PP | 6.1 | 32.2 | 3.6 | 0.002 |
| RH | 4.5 | 23.7 | 2.7 | 0.002 |
| RLAI | 2.9 | 15.1 | 1.8 | 0.012 |
| AT | 2.9 | 15.2 | 1.8 | 0.022 |
| S | 2.6 | 13.9 | 1.7 | 0.020 |

## Discussion

### Bryophyte heterogeneity

Bryophytes studies in pure and mixed forests in southern Patagonia are very lack even considering the high mosses or liverwort diversity [18, 19]. Here, we observed that microhabitats did not differ greatly between forest types and landscape. Diversity vary between forest types (similar between mixed and evergreen), as well as similar between landscape. Evergreen forests, where bryophytes were more diverse than other forests, and mixed forests were found on average between both pure deciduous and evergreen forests.

Here, we reported the mixed forest had more exclusive species than pure deciduous or evergreen forests. Of these species, only one liverwort (*Clasmatocolea sp.*) was found in both landscape types, but some mosses (*Bartramia mossmaniana*, *Hymenodontopsis mnioides*, *Hypnum skottsbergii*) and the liverwort (*Lophozia sp.*) and were found only on the coast. Therefore, the mixed *Nothofagus* forests had unique conditions capable of developing exclusive species that are not in pure forests.

As mentioned above, the role of the diverse microclimate and microhabitat would be directly related to the indicator species [1, 36, 37]. We found two mosses (*Dicranoloma robustum* and *Ditrichum cylindricarpum*) that were indicative of evergreen forests and they were growing on litter, decaying wood or as epiphytes microsites. Notably, some species were indicator of forest types and landscape types, being remarkable for conservation strategies. For example, the moss (*Campylopus clavatus*) was indicator of mixed forests, and the moss (*Acrocladium auriculatum*) was indicator of deciduous forests. Therefore, this remarked the importance of microhabitat quality for bryophytes.

The bryophyte communities in *Nothofagus* forests could be considered endemic and specific to these southern areas. The pattern of distribution of bryophyte species in the forests could help to understand how species were distributed according to climatic conditions and microhabitats. For example, León et al. [30] indicated that it is important to know the patterns of endemic species in relation to environmental preferences. In our results, the origin of 42% bryophytes is the southern Antarctic for deciduous forests. This coincides with studies of bryophytes in endemic Valdivian forests of South America, where Villagrán et al. [32] suggests that there is prolonged isolation of the species.

### Environmental drivers and influence of canopy layer composition in coast and mountain

Our study showed that bryophyte communities were significantly related to the microclimate (particularly the effective precipitation, relative air humidity, and air temperature), besides to the slope and the RLAI, which would be determined by the influence of the canopy composition and climate in the landscape type. Our results were in concordance with previous studies in the northern hemisphere [1, 38, 39], which indicates the bryophyte community composition is influenced by the combination of microclimate conditions, which may vary according to the forest community in which they are found.

Specific environmental drivers could influence the richness, cover and diversity of bryophytes. In our study, air conditions seem to be highly important for bryophyte composition in the two gradients under analysis (canopy composition and landscape), where differences were important for each forest type and landscape type. Precipitation was also crucial to water storage in soils and thus keeping environmental moisture in the forest sites. However, the water storage capacity in these forest soils vary greatly, and generates differences in the soil moisture, and therefore differences in the bryophyte abundance as suggested by Raabe et al. [40]. In this context, the regional differences (altitudinal gradients) for the cover, richness and diversity of bryophyte species are evidently determined by climatic differences, like relative air humidity and soil moisture, but also geological drivers such as landscape texture and soil types, which offer a wide range of microhabitats for bryophytes [38]. Here, the landscape types had a strong effect on the soil physical-chemical conditions differing noticeably between the coast and the mountain [14]. It was observed that the slope differentiates the coast of the mountain, mainly due to the altitude in which the forest types develop. In addition, air temperature is considered a key driver structuring the bryophyte community in different forest types [40]. Variations in temperature, together with higher relative air humidity were considered favourable conditions for bryophytes abundance [38]. However, high temperatures were also considered to limit the abundance of bryophytes in the forest [41]. Notably, this can be one reason for the differences between the bryophytes between the coast and mountain because the coast had higher temperatures generating greater drying of the soils in forest in the coast. Contrary, mountain soils had greater moisture (air and soil) maintaining favourable microclimatic conditions for bryophytes development.

Previous studies indicate that the availability of atmospheric water is more important for bryophytes than the level of soil moisture since bryophytes absorb water throughout their entire plant structure [42]. Therefore, in forest ecosystems a high level of air humidity surrounding the understorey is important for the bryophytes, as well as shaded conditions [43]. Our study showed that the relative air humidity was higher on the coast (influenced by the proximity to the sea), this driver was the one that probably best influences the composition of bryophytes (mainly in mixed forests and evergreen forests), while the soil moisture explains better the specific composition in the mountains. Raabe et al. [40] considered that soil moisture is the main driver of bryophyte diversity. In our study, although the higher soil moisture in mountain forests presented a greater coverage of bryophytes, the less humid soils on the coast for the three forest types could be important to conserve rare species, even if they are present in low abundance.

Other specific drivers can also influence richness, cover and diversity of bryophytes along with the microclimate. Changes in the forest canopy cover (i.e., forest structure, RLAI, etc.) mainly in the deciduous forests along the year (e.g., complete foliage development in summer and loss of 50% foliage in autumn-winter), determine high variation in the solar radiation reaching the ground, wind exposition, and air and soil moisture [3]. All these drivers are less variable within mixed and the evergreen forest where more than half of the canopy remain green and developed year round. Tinya et al. [44] found that radiation correlates positively with the bryophytes richness on the forest floor, although direct exposure of radiation can also cause desiccation. However, in our study the solar radiation was not considered as main driver of richness and cover of bryophytes. In coniferous forests, bryophytes richness are benefited from greater radiation availability in the forest floor due to a less closed canopy and a lower litter drop compared to deciduous forests [44]. This could be a point to consider in mixed and evergreen forests of south Patagonia during autumn and spring when the foliage of deciduous trees is still incomplete. However, we found there were not major differences in canopy cover during summer, so it can occur in very specific seasons or places (e.g., in gaps), which was not considered in this study but it is an interesting aspect of the functioning of these mixed forests to consider in future researches.

The study by Marialigeti et al. [45] indicated the litter of deciduous species changes the properties of soils, which can inhibit the presence of bryophytes. We have observed that bare soil with litter is the microhabitat with the largest number of liverworts and mosses in all forest types and landscape types. In the study area, Toro Manríquez et al. [14] calculated that the litterfall for deciduous forest exceeds in 26.4% to the mixed forest and 51% to the evergreen forest. Therefore, this microenvironmental driver is essential for the richness, cover and diversity of species. According to Müller et al. [38], the presence of different substrates in beech forests of central Europe can better favour vascular plants, but are less suitable for bryophytes occurrence. This could explain the lower richness, cover and diversity of bryophytes in deciduous forests, which have greater vascular plants cover and lichens cover, than mixed and evergreen forests.

Previous studies showed that the presence of bare soil and in places with slopes (where occasional soil disturbances can occur) hinder the growth of bryophytes [38, 45]. Although we did not recorded the occurrence of bryophytes in relation to slopes, they grew on bare soil mainly in the deciduous and mixed forests in the coast and mountain. Probably, this substrate was more used by mosses and liverworts because are less suitability for vascular plants (e.g., shrubs, forbs). Müller et al. [38] found that the richness of bryophytes in the forest floor decreases with the increase in soil pH. In our study, evergreen forests had lower pH values than mixed and deciduous forests. This could be interesting to analyze considering the substrates in which liverworts and moss grow.

Decaying wood is considered an important substrate mainly for the bryophytes cover [39, 40, 43, 46] since this substrate retains moisture providing a more stable microclimate at soil level [7]. Rajandu et al. [47] suggested liverworts species depends mainly on wetter microhabitats such as decomposing wood. In our study, the richness and cover of liverworts were greater in evergreen forests, and the microhabitat of decaying wood was essential for bryophytes occurrence in that forest type.

The bryophytes epiphytes on branches and bark are incorporated to ground-bryophyte communities after the fall of tree structures. Therefore, some mosses and liverworts develop as epiphytes on some part of *Nothofagus* trees and then in the soil, so they remain and continue their development as long as the appropriate conditions. This is important to consider in conservation strategies because the heterogeneity of old-growth forests contribute to the microhabitat diversity where bryophytes can occur [39, 45].

The occurrence of suitable microhabitat for wide variety of bryophytes can generate favourable conditions for the ecological functioning of the forest (e.g., seedling regeneration) [38, 40]. Rehm et al. [39] indicated that bryophytes could be better substrate for regeneration seedlings than leaf litter, and even as good as decaying wood. In addition, one of the properties that bryophytes have on seedlings is the ability to retain seeds, regulate soil moisture and not compete with regeneration plants such as other vascular plants [48]. However, our results showed not all bryophytes act as a substrate for seedlings in the different forest types, (i.e., in mixed and evergreen forests, liverwort or mosses were not related to seedlings). Nevertheless, when the landscape types were considered, bryophytes seem to be a relevant substrate for *Nothofagus* seedlings more important in the mountain than in the coast.

### Implications for conservation and general considerations

Our study indicates microhabitat was a fundamental characteristic to maintain the diversity of bryophytes. In previous studies, the association and facilitation offered by bryophytes to regeneration seedlings was considered as a measure in restoration [39, 48, 49]. The study of the life forms of the bryophyte species and the association of the communities must be considered in the future. These associations respond to the different microclimatic conditions of the forest in particular (e.g., radiation tolerance, soil moisture, microhabitat, seed density and forest regeneration).

Our results reinforce the idea that hat *Nothofagus* forests not configured continuous ecological elements from sea level to the treeline [50, 51]. The mixed forests showed particularities such as the appearance of species that are not found in pure stands, showing that different forest types show a particular diversity. In this study, the forests in the coast were located within a protected area (Tierra del Fuego National Park), so this information is valuable to spread knowledge about conspicuous and little-known plant species and particularly for bryophyte conservation. Cold-adapted mountain species are generally sensitive to climate change [52, 53]. In the mountain area, conservation actions focused on bryophyte community must consider some of the aspects studied here such as air temperature and humidity, precipitation or soil conditions to face future climate change. Due to different species of bryophytes had different temperature sensitivities, each species must react differently to potential climate change Previous studies on different biological groups affirm that species composition of terrestrial ecosystems will be modified with the expected global change (e.g., temperature increase, drought) [54, 55]. Indeed, in the face of the climate change scenario and with the increase in global radiation, a decrease in the abundance of bryophytes is expected [40]. Therefore, our results highlight the importance of understands the environmental drivers affecting the richness and cover of bryophytes. This could be valuable information for conservation of bryophyte species, and to generate basic knowledge about the potential effects of climate change in this area.

## Conclusions

The forest types and landscape types influence differentially the richness, cover and diversity of bryophytes communities. This was conditioned by the microclimate into the forests (i.e., soil moisture, relative air moisture and air temperature), as well as by characteristics of the regional climate influenced by the landscape type. The diversity of microhabitats was essential for mosses and liverworts development, such as those generated by litter and decaying wood. Here, the evergreen forest presented greater cover of most favorable microhabitats along with the mixed forests. However, the mixed forest presenting exclusive species of bryophytes absents in deciduous and evergreen forests (mainly in the coast landscape). Therefore, the mixed forest represented an intermediate situation but rather a particular one influenced by the combination of deciduous and evergreen proportion of canopies. On the other hand, the mountain sustains the greater diversity and cover of bryophytes in all forest types. According to distribution patterns, bryophytes of *Nothofagus* forests could be considered endemic and highly specific to these southern areas. This study constitutes an important basis for continuing studies of bryophytes in the southernmost forests, investigating their role in forest dynamics and as a key component of local and regional biodiversity. Conservation efforts should not only consider the forest type, but also the landscape type and therefore the microclimatic conditions. Our study highlights the crucial effect of climatic conditions on the composition of bryophyte community, thus this group could be more sensitive to the expected changes of temperature and precipitations. This knowledge could improve ground-bryophyte species conservation at stand and landscape level in temperate forests.

## Acknowledgements

We thank the Centro Austral de Investigaciones Científicas (Ushuaia, Argentina), Museo Nacional de Historia Natural (Santiago de Chile, Chile) and Tierra del Fuego National Park (Ushuaia, Argentina) for their support during the realization of this study.

## Contributor role

Conceptualization = Mónica Toro Manríquez

Data Curation = Mónica Toro Manríquez, Víctor Ardiles

Formal Analysis = Mónica Toro Manríquez, Álvaro Promis, María Vanessa Lencinas

Investigation = Mónica Toro Manríquez, Rosina Soler

Methodology = Mónica Toro Manríquez, Víctor Ardiles, Álvaro Promis, Maria Vanessa Lencinas

Project Administration = Mónica Toro Manríquez

Resources = Víctor Ardiles, María Vanessa Lencinas, Guillermo Martínez Pastur

Supervision = Álvaro Promis, Guillermo Martínez Pastur

Validation = Víctor Ardiles

Writing – Original Draft = Mónica Toro Manríquez, Alejandro Huertas Herrera

Writing – Review & Editing= Mónica Toro Manríquez, Alvaro Promis, Alejandro Huertas Herrera, Rosina Soler, María Vanessa Lencinas, Guillermo Martínez Pastur

## Supporting information

**S1 Table. Bryophyte species observed in each forest type** (Np = deciduous forests with >80% *Nothofagus pumilio* canopy cover, M = mixed forests of *N. pumilio* and *N. betuloides*, Nb = pure evergreen forests with >80% *N. betuloides* canopy cover) and landscape types (COA = coast, MOU = mountain), showing: (i) species code, (ii) TAX= taxonomic group (Li = liverworts, Ms = mosses), (iii) global distribution patterns-GDP (D = disjunt with South America, South Africa and Europe, E = endemic, PAN = pantropical-type *Podocarpus*, A = austral-antarctic, COS = cosmopolitan, B = bipolar), and (iv) microhabitat or substrate (LT = litter cover, DW = decaying wood, BS = bare soil, St = stone, EP = epiphytic on branches and bark in the forest floor). OF = occurrence frequency in each forest type (%), and ẋ = mean frequency of occurrence in the entire study (%).

**S2 Table. Two-way ANOVAs evaluating the effect of tree species contribution in the canopy composition in mixed *Nothofagus* stands and landscape types (COA = coasts, MOU = mountains) over the forest structure variables:** BA = basal area (m^2^ ha^−1^), DH = dominant height (m), DBH = diameter at breast height (cm).

*F, p* = Fisher test and probability. Different letters in each column show significant differences by Tukey test at p <0.05

**S3 Table. Crosstabs of frequency and chi-square test of liverworts and mosses for each microhabitat.** LT = litter, DW = decaying woods, BS = bare soil, St = stone, EP = epiphyte of branches and bark in the forest floor. CNp= deciduous *Nothofagus pumilio* forests in the coasts, CM= mixed *N. pumilio* and *N. betuloides* forests in the coasts, CNb = evergreen *N. betuloides* forests in the coasts, MNp = deciduous *N. pumilio* forests in the mountains, MM= mixed *N. pumilio* and *N. betuloides* forests in the mountains, MNb = pure evergreen *N. betuloides* forests in the mountains.

**S4 Table. Crosstabs of frequency and chi-square test of tree seedlings found growing jointly with bryophytes (liverworts and mosses) for each forest type and in each landscape types. Species code are presented in S1Table.** CNp= deciduous *Nothofagus pumilio* forests in the coasts, CM= mixed *N. pumilio* and *N. betuloides* forests in the coasts, CNb = evergreen *N. betuloides* forests in the coasts, MNp = deciduous *N. pumilio* forests in the mountains, MM= mixed *N. pumilio* and *N. betuloides* forests in the mountains, MNb = pure evergreen *N. betuloides* forests in the mountains.

**S5 Table. Pearson correlation coefficients between variables tested for Canonical Correspondence Analysis.** BA = basal area (m^2^ ha^−1^), DH = dominant height (m), DBH = diameter at breast height (cm), CC = canopy cover (%), RLAI = relative leaf area index, GR = transmitted global radiation (%), SM soil moisture (%), PP = annual precipitation (mm yr^−1^), ST = soil temperature (°C), AT = air temperature (°C), RH = relative air humidity (%), S = slope (%), pH, R = resistance to penetration (N m^−2^), BS = bare soil cover (%), Ds = debris cover (%), VC = vascular plant cover including ferns, monocots and dicots (%), and L = lichen cover (%).

**S1 Fig. Cover (%) of bryophyte species for each microhabitat.** (A) liverworts, (B) mosses, analysing LT = litter cover, DW = decaying wood, BS = bare soil, EP = epiphytic on branches and bark in the forest floor, St = stone. Species code are presented in S1Table.

## References

1. Jiang T, Yang X, Zhong Y, Tang Q, Liu, Su Z. Species composition and diversity of ground bryophytes across a forest edge-to-interior gradient. Sci Rep. 2018;8: 11868. doi: 10.1038/s41598-018-30400-1.

2. Vanderpoorten A, Goffinet B. Introduction to bryophytes. 1st ed. Cambridge, UK: Cambridge University Press; 2009.

3. Bartels SF, Macdonals SE, Johnson D, Caners RT, Spence JR. Bryophyte abundance, diversity and composition after retention harvest in boreal mixedwood forest. J Appl Ecol. 2018;55: 947–957. doi: 10.1111/1365-2664.12999.

4. Jeschke M, Kiehl K. Effects of a dense moss layer on germination and establishment of vascular plants in newly created calcareous grasslands. Flora. 2008; 203: 557–566. doi:10.1016/j.flora.2007.09.006.

5. Delach A, Kimmerer RW. The effect of *Polytrichum piliferum* on seed germination and establishment on iron mine tailings in New York. Bryologist. 2002;105: 249–255. doi:10.1639/0007-2745(2002)105[0249:TEOPPO]2.0.CO;2.

6. Sun SQ, Wu YH, Wang GX, Zhou J, Yu D, Bing HJ, Luo J. Bryophyte Species Richness and Composition along an altitudinal gradient in Gongga mountain, China. PLoS One. 2013; 8: 3: e58131. doi: 10.1371/journal.pone.0058131.

7. Mills SE, Macdonald SE. Predictors of moss and liverwort species diversity of microsites in conifer-dominated boreal forest. J Veg Sci. 2004;15: 189–198. doi:10.1658/1100-9233(2004)015[0189:POMALS]2.0.CO;2.

8. Mills SE, Macdonald SE. Factors influencing bryophyte assemblage at different scales in the western Canadian boreal forest. Bryologist. 2005; 108: 86–100. doi:10.1639/0007-2745(2005)108[86:FIBAAD]2.0.CO;2.

9. Márialigeti S, Tinya F, Bidló A, Ódor P. Environmental drivers of the composition and diversity of the herb layer in mixed temperate forests in Hungary. Plant Ecol. 2016;217: 549–563. doi:10.1007/s11258-016-0599-4.

10. Tinya F, Ódor P. Congruence of the spatial pattern of light and understory vegetation in an old - growth, temperate mixed forest. For Ecol Manage. 2016;381: 84–92. doi:10.1016/j.foreco.2016.09.027.

11. Frahm J-P, Vitt DH. A comparison of the mossfloras of Europe and North America. Nova Hedwigia. 1993;56: 307–333.

12. Frahm J-P. Diversity, dispersal and biogeography of bryophytes (mosses). Biodivers Conserv. 2008; 17: 277–284. doi:10.1007/s10531-007-9251-x.

13. Toro Manríquez M, Mestre L, Lencinas MV, Promis Á, Martínez Pastur G, Soler R. Flowering and seeding patterns in pure and mixed *Nothofagus* forests in southern Patagonia. Ecol Process. 2016;5: 21–33. https://doi.org/10.1186/s13717-016-0065-1.

14. Toro Manríquez M, Soler R, Lencinas MV, Promis A. Canopy composition and site are indicative of mineral soil conditions in Patagonian mixed *Nothofagus* forests. Ann For Sci. 2019;76: 117. doi:10.1007/s13595-019-0886-z.

15. Mestre L, Toro-Manríquez M, Soler R, Huertas-Herrera A, Martínez Pastur G, Lencinas MV. The influence of canopy-layer composition on understory plant diversity in southern temperate forests. For Ecosys. 2017;4: 6. doi:10.1186/s40663-017-0093-z.

16. Ah-Peng C, Chuah-Petiot M, Descamps-Julien B, Bardat J, Stamenoff P, Strasberg D. Bryophyte diversity and distribution along an altitudinal gradient on a lava flow in La Réunion. Divers Distrib. 2007;13: 654–662.

17. Gignac LD. Bryophytes as indicators of climate change. Bryologist. 2001; 104: 410–420.

18. Moore D. Flora of Tierra del Fuego. London: Anthony Nelson; 1983.

19. Greene SW, Hässel de Menéndez GG, Matteri CM. La contribución de las briófitas en la vegetación de la transecta. In: Boelcke O, Moore DM, Roig FA, editors. Transecta Botánica de la Patagonia Austral. Argentina: Consejo Nacional de Investigaciones Científicas y Técnicas Chile: Instituto de la Patagonia UK: Royal Society; 1985. pp. 557–591.

20. Lencinas MV, Martínez Pastur G, Solán R, Gallo E, Cellini JM. Forest management with variable retention impact over bryophyte communities of *Nothofagus pumilio* understory. Forstarchiv. 2008; 79: 77–82.

21. Promis A, Gärtner S, Reif A, Cruz G. Effects of canopy gaps on forest floor vascular and non-vascular plant species composition and diversity in an uneven-aged *Nothofagus betuloides* forest in Tierra del Fuego, Chile. Community Ecol. 2012; 13:2: 145–154. doi:10.1556/comec.13.2012.2.3.

22. Contreras H, Borgel R, Quezada M, García de Cortázar V, Rojas M, Bitterlich W. Informe de la primera etapa del proyecto sobre reforestación de la Precordillera Patagónica (Cuadrángulos Skyring y Rubens). Facultad de Ciencias Forestales, Santiago: Universidad de Chile Press; 1975.

23. Bitterlich W. The relascope idea. Relative measurements in forestry. London: CAB Press; 1984.

24. Frazer GW, Fournier RA, Trofymow JA, Gall RJ. A comparison of digital and film fisheye photography for analysis of forest canopy structure and gap light transmission. Agric For Meteorol. 2001;109: 249–263.

25. Levy EG, Madden EA. The point method of pasture analyses. N Z J Agric 1993;46: 267–379.

26. Ardiles V, Peñaloza A. Briófitas del área urbana de Santiago de Chile: Especies, hábitats y consideraciones para su conservación. Boletín del Museo Nacional de Historia Natural. 2013; 62: 95–117.

27. Ardiles V, Cuvertino J, Osorio YF. Briófitas de los bosques templados de Chile. Una introducción al mundo de los Musgos, Hepáticas y Antocerotes. Guía campo. 1st ed. Santiago, Chile: CORMA Press; 2008.

28. Müller F. An updated checklist of the mosses of Chile. Archive for Bryology. 2009;58: 1–124.

29. Hässel de Menéndez GG, Rubies MF. Catalogue of the Marchantiophyta and Anthocerotophyta of southern South America. Chile, Argentina and Uruguay, including Easter Is. (Pascua I.), Malvinas Is. (Falkland Is.), South Georgia Is., and the subantarctic South Shetland Is., South Sandwich Is., and South Orkney Is. Nova Hedwigia. 2009;134:1–672.

30. León CA, Oliván G, Larraín J, Vargas R, Fuertes E. Bryophytes and lichens in peatlands and *Tepualia stipularis* swamp forests of Isla Grande de Chilé (Chile). An Jard Bot Madr. 2014;71: 1. doi:10.3989/ajbm.2342.

31. Seki T. A moss flora of Provincia de Aisén, Chile. Journal of Science of the Hiroshima University, Series B, Div. 2 (Botany). 1974;15: 9–101.

32. Villagrán C, Hässel de Menéndez G. Barrera E. Hepáticas y Anthocerotes del Archipiélago de Chiloé. Una introducción a la flora briofítica de los ecosistemas templados lluviosos del sur de Chile. Santiago de Chile: Corporación de Amigos del Museo Nacional de Historia Natural Press; 2005.

33. Pielou EC. Mathematical Ecology. New York: John Wiley and Sons Inc. Press; 1975.

34. Dufrêne M, Legendre P Species assemblages and indicator species: the need for a flexible asymmetrical approach. Ecol Monog. 1997;67: 345–366.

35. McCune B, Mefford MJ. Multivariate analysis of ecological data, Version 4.0, MjM software. Oregon: Gleneden Beach Press;1999.

36. Fenton NJ, Bergeron Y. Does time or habitat make old-growth forests species rich? Bryophyte richness in boreal *Picea mariana* forests. Biol Conserv. 2008;141: 1389–1399. doi:10.1016/j.biocon.2008.03.019.

37. Boudreault C, Paquette M, Fenton NJ, Pothier D, Bergeron Y. Changes in bryophytes assemblages along a chronosequence in eastern boreal forest of Quebec. Can J For Res. 2018; 48:7: 821–834. doi.10.1139/cjfr-2017-0352.

38. Müller J, Boch S, Pratic D, Socher DA, Pommerag U, Hessenmöller D. et al. Effects of forest management on bryophyte species richness in Central European forests. For Ecol Manage. 2019;432: 15: 850–859. doi:10.1016/j.foreco.2018.10.019.

39. Rehm EM, Thomas MK, Yelenik SG, Bouck DL, D’ Antonio CM. Bryophyte abundance, composition and importance to woody plant recruitment in natural and restoration forests. For Ecol Manage. 2019;444: 405–413. doi:10.1016/j.foreco.2019.04.055.

40. Raabe S, Müller J, Manthey M, Dürhammerd O, Teuber U, Göttlein A. et al. Drivers of bryophyte diversity allow implications for forest management with a focus on climate change. For Ecol Manage. 2010;260: 11:1956–1964. doi.10.1016/j.foreco.2010.08.042.

41. Berdugo MB, Quant JM, Wason JW, Dovciak, M. Latitudinal patterns and environmental drivers of moss layer cover in extratropical forests. Glob Ecol Biogeogr. 2018;27: 10: 1213–1224. doi.org/10.1111/geb.12778.

42. Graf MD, Rochefort L. Moss Regeneration for Fen Restoration: Field and Greenhouse Experiments. Restor Ecol. 2010;18: 1:121–130. doi:10.1111/j.1526-100X.2008.00437.x-.

43. Ódor P, Király I, Tinya F, Bortignon F, Nascimbene J. Patterns and drivers of species composition of epiphytic bryophytes and lichens in managed temperate forests For Ecol Manage. 2013;306: 256–265. doi:10.1016/j.foreco.2013.07.001.

44. Tinya F, Márialigeti S, Király I, Németh B, Ódor P. The effect of light conditions on herbs, bryophytes and seedlings of temperate mixed forests in Orség, Western Hungary Plant Ecol. 2009;204: 1: 69–81. doi: 10.1007/s11258-008-9566-z.

45. Márialigeti S, Németh B, Tinya F, Ódor P. The effects of stand structure on ground-floor bryophyte assemblages in temperate mixed forests. Biodivers Conserv. 2009; 18: 2223–2241. doi:10.1007/s10531-009-9586-6.

46. Rudolphi J, Gustafsson L. Forests regenerating after clear-cutting function as habitat for bryophyte and lichen species of conservation concern. PLoS ONE. 2011; 6: 4: e18639. doi:10.1371/journal.pone.0018639.

47. Rajandu E, Kikas K, Paal J. Bryophytes and decaying wood in *Hepatica* site-type boreo-nemoral *Pinus sylvestris* forests in Southern Estonia. For Ecol Manage. 2009; 257: 3: 994–1003. doi.org/10.1016/j.foreco.2008.11.001.

48. Inman-Narahari F, Ostertag R, Asner GP, Cordell S, Hubbell SP, Sack L. Trade-offs in seedling growth and survival within and across tropical forest microhabitats. Ecol Evol. 2014; 4: 19: 3755–3767. doi: 10.1002/ece3.1196.

49. Michalska-Hejduk D, Wolski GJ, Harnisch M, Otte A, Bomanowska A, Donath TW. Restoration of floodplain meadows: Effects on the re-establishment of mosses. PLoS ONE. 2017; 12:12:e0187944. doi:10.1371/journal.pone.0187944.

50. Martínez Pastur G, Peri PL, Soler RM, Schindler S, Lencinas MV. Biodiversity potential of *Nothofagus* forests in Tierra del Fuego (Argentina): Tool proposal for regional conservation planning. Biodiv. Conserv. 2016;25: 1843–1862. doi:10.1007/s10531-016-1162-2.

51. Huertas Herrera A, Cellini J.M, Barrera MD, Lencinas MV, Martínez Pastur GJ. Environment and anthropogenic impacts as main drivers of plant assemblages in forest mountain landscapes of Southern Patagonia. For. Ecol. Manage. 2018; 430: 380–393. doi:10.1016/j.foreco.2018.08.033.

52. Bergamini A, Ungricht S, Hofmann H. An elevational shift of cryophilous bryophytes in the last century - an effect of climate warming? Divers Distrib. 2009; 15: 871–879. doi:10.1111/j.1472-4642.2009.00595.x.

53. Thuiller W, Lavorel S, Araujo MB, Sykes MT, Prentice IC Climate change threats to plant diversity in Europe. Proc Natl Acad Sci USA; 2005;102: 23: 8245–8250. doi:10.1073/pnas.0409902102.

54. Kelly A, Goulden M. 2008. Rapid shifts in plant distribution with recent climate change. Proc Natl Acad Sci USA. 2008; 105: 11823–11826. doi:10.1073/pnas.0802891105.

55. Lenoir J, Gégout JC, Marquet PA, De Ruffray P, Brisne H. A significant upward shift in plants species optimum elevation during the 20^th^ century. Science. 2008;320: 1768–1771. doi:10.1126/science.1156831.

